# Performance of *Bracon brevicornis* (Wesmael) on two *Spodoptera* species and application as potential biocontrol agent against fall armyworm

**DOI:** 10.1101/2020.06.27.171025

**Authors:** Enakshi Ghosh, Richa Varshney, Radhika Venkatesan

## Abstract

Successful pest management using parasitoids requires careful evaluation of host-parasitoid interactions. Here, we report the performance of larval ecto-parasitoid wasp, *Bracon brevicornis* (Wesmael) on important agricultural pests, *Spodoptera litura* (Fabricius) and *S. frugiperda* (J.E. Smith). Biology of *B. brevicornis* was studied on different host instars under laboratory and cage setup. In no-choice assay, the parasitoid development was highest on fifth instar *S. litura* larvae as the wasp laid ∼253 eggs with 62% hatching, 76% pupae formation and 78% adult emergence. Similarly, these parameters were highest on fifth instar *S. frugiperda* larvae (293 eggs, 57% hatching, 80% pupae formation, 70% adult emergence). In two-choice assay, *B. brevicornis* preferred fourth or fifth over third instar larvae of both hosts. Successful parasitism depends on host paralysis and suppression of host immunity. *B. brevicornis* interaction downregulated cellular immunity of both hosts as shown by reduced hemocyte viability and spreading. The percent parasitism rate of *B. brevicornis* was unaltered in the presence of host plant, *Zea mays* in cage study. 76 and 84% parasitism was observed on fifth instar larvae of *S. litura* and *S. frugiperda*, respectively. We evaluated the performance of *B. brevicornis* as a biocontrol agent on *S. frugiperda* in maize field. Our results show 54% average reduction in infestation after release of *B. brevicornis*. Taken together, we report the performance of *B. brevicornis* on important insect pests for the first time in laboratory and field conditions. Our findings indicate that *B. brevicornis* is a promising candidate for integrated pest management.

**Key messages:**

1. We have evaluated the instar preference and performance of *B. brevicornis* as a potential biocontrol agent for two serious pests, *Spodoptera litura* and *S. frugiperda*.
2. Fifth instar larva was most suitable for *B. brevicornis* development irrespective of the host species. *B. brevicornis* attack induced permanent paralysis and down-regulated cellular immunity of both hosts.
3. Our field experiment confirmed *B. brevicornis* as a promising parasitoid for controlling *S. frugiperda*, a highly invasive pest of growing concern.

## Introduction

Insect parasitoids are fascinating model systems to study ecological interactions and are of high economic importance as biological control agents. As adults, parasitoids are free-living whereas in the larval phase, they need to feed on a single host to reach adulthood. For this, they develop either inside or on the host after inducing temporary or permanent paralysis by the adult parasitoid. At the end of their feeding, the hosts are killed (Godfray 1994). Ectoparasitoids inject maternal factors that cause permanent paralysis before oviposition making them effective natural enemies for pest management (Qian et al. 2013; Quicke 2015). *Bracon brevicornis* (Wesmael) (Hymenoptera: Braconidae) is a gregarious parasitoid that is known to attack stored grain and crop pests. They are idiobionts and develop concealed or semi-concealed on larval hosts (Quicke 2015). Females are synovigenic as their eggs mature throughout their lifetime and they exhibit host feeding behavior to acquire nutrients. Females also sometimes paralyze and host feed without depositing any eggs. *Bracon brevicornis* has been recorded on several host species but detailed study of their biology on *Spodoptera* spp.is lacking and is needed for proper utilization in management programs (Dabhi et al. 2011).

The Noctuid genus *Spodoptera* (Lepidoptera) encompasses 31 species, half of which are considered as serious pests worldwide, commonly referred to as ‘armyworms’ (Pogue 2002). Among these, *Spodoptera litura* (Fabricius) and *Spodoptera frugiperda* (J.E. Smith) are important pest species that are highly polyphagous. *Spodoptera litura*, widely distributed throughout tropical and subtropical regions is a notorious pest of more than hundred plant species. For instance, in India, the economic impact of *S. litura* ranges between 30-98% of major crops like soybean and groundnut (Dhir 1992; Natikar 2015). In addition to high fecundity, short life cycle and polyphagy, resistance to chemical as well as biopesticides (Bt) is another concern (Cheng et al. 2017). The fall armyworm, *S. frugiperda* (FAW) is native to North America and it was first recorded in African continent in 2016 (Day et al. 2017; Goergen et al. 2016), then subsequently spread very quickly across Asia causing severe damage to cereal crops (Sharanabasappa 2018). In India, it was first reported in 2018 on Maize and has now spread to 20 states (Shylesha et al. 2018). It is known to attack at least 80 different crop plants with preference towards cereals, especially maize. In African countries, the value of these losses are estimated to be ∼2.48 -6.19 billion USD (Prasanna et al. 2018). Effective control of FAW is an urgent need as the impact of FAW infestation on maize yield is alarming and favorable agro-ecological conditions can lead to large pest outbreaks causing lasting threat to important crops (Goergen et al. 2016). Natural enemies of insect pests exert top-down pressure and control insect populations. Despite reports on natural enemies, very few studies report larval parasitoids and their potential in controlling FAW (Sisay et al. 2018).

The success of parasitoid depends on various factors related to their host *viz*. developmental stage, size, diet as well as environmental conditions (Pan et al. 2017; Khan et al. 2016). Studies comparing parasitoid performance and host size have shown positive correlation as these factors can directly influence parasitoid fitness (Charnov 1982; Stoepler et al. 2011). Similarly, host plant quality and chemistry also influence parasitoid behavior, development and fitness (Gols et al. 2008; Sarfraz et al. 2019). Evaluation of these parameters is crucial to understand parasitoid biology, which in turn, is important for field application. Here, a comprehensive study on *B. brevicornis* performance on two lepidopteran hosts; *S. litura* and *S. frugiperda* is presented, with potential application for biological control in field conditions. Together, our results show that *B. brevicornis* is efficient in controlling FAW and could play an important role in integrated pest management programs of FAW.

## MATERIALS AND METHODS

### Insect Rearing Conditions

Laboratory culture of *S. litura* (NBAII-MP-NOC-02) and *S. frugiperda* (NBAIR-MP-NOC-03) were reared on castor bean plant (*Ricinus communis*) in a climatic chamber. *Bracon brevicornis* was collected from Coimbatore, India and maintained on laboratory host, the rice moth *Corcyra cephalonica* (Stainton) (Srinivasan and Chandrikamohan 2017). The adult wasps were provided with 50% honey-water solution (v/v). 3-day-old mated female wasps were used for our study. Rearing of insects and all laboratory experiments were conducted at 26°C with a photoperiod of 12L:12D and 60% relative humidity.

### Biology of *B. brevicornis* on different host instars

*B. brevicornis* is known to parasitize late instars of *S. litura* larvae (Ghosh and Venkatesan 2019). Hence, 3-5^th^ instar host larvae were chosen for this study. We performed no choice and two choice tests to assess the preference and performance of *B. brevicornis*. For no-choice assay, in a glass test tube (measuring 160mm x 16mm x 14mm), single larva (of a specific instar) of *S. litura* or *S. frugiperda* was kept with one mated female and male of *B. brevicornis* for 24h. Post 24h, the wasp couple was removed and placed into another test tube with a fresh larva of same instar for parasitization. The adult wasps were provided with 50% honey-water solution (v/v). This experiment was repeated until the death of the particular female wasp and replicated at least 5 times independently. The parasitized larvae were kept under same conditions and the total number of eggs laid (life-time fecundity/female wasp), percent larval hatching (number of larvae/total number of eggs laid x 100), percent pupae formation (number of pupae/total number of larvae x 100), adult emergence and number of females were recorded for comparative analysis. For calculating the fecundity, total number of eggs laid by the parasitoid was manually counted under a stereoscope. For two choice assay, we performed five sets of experiments. In each set, one female of *B. brevicornis* was released in a petri-plate and given a choice of: (a) third vs fourth, (b) third vs fifth and (c) fourth vs fifth instar larvae of *S. litura* and *S. frugiperda*, separately. Five female parasitoids were tested for their choice in each set. Single female wasp was released in the middle of the petri-plate and observed for a maximum duration of 6h or until at least one larva was paralyzed.

### Biology of *B. brevicornis* in the presence of host plant

Parasitoids utilize plant volatiles as cues to locate their host and several studies have shown the impact of host plant in modulating parasitoid behavior (Stoepler et al. 2011; Li et al. 2014; Girling et al. 2011; Kugimiya et al. 2010). As *S. frugiperda* is serious pest on maize, we recorded the biology of *B. brevicornis* in the presence of maize. For this, plants (*Zea mays* DHM103) were grown in a pot (15 × 10 cm) with red soil and coco peat in green house (26±5°C) under natural light conditions. No insecticide was sprayed on plants during experiment. Twenty-days old maize plants were used for the study. In a cage measuring 60 × 50 cm, five potted maize plants were arranged in a randomized block design. *S. litura* or *S. frugiperda* larvae of specific instar (3^rd^, 4^th^ or 5^th^) were placed on plants in 1: 3 ratios (plant: larvae) and allowed to feed for 24h. Then, 3-day old mated females and males of *B. brevicornis* were released in 1: 3 ratio (wasp: larvae) into the cage and allowed to interact for 24h. The instars of *S. litura* and *S. frugiperda* were tested separately as independent experiments. Percent parasitization was recorded considering the number of paralyzed larvae. The parasitized larvae from each treatment were kept in a test tube (160mm x 16mm x 14mm) until adult emergence. The number of eggs found on each larva, the percent adult emergence and their longevity were recorded.

### Regulation of host immune-competence by *B. brevicornis*

Parasitoid venom is known to suppress host immune functions that help in better survival of their progeny (Teng et al. 2016; Ghosh and Venkatesan 2019). This includes reduction in hemocyte viability and spreading ability that are important to combat infections. Castor fed fifth instar larva of *S. litura* and *S. frugiperda* was placed in a petri dish along with one mated female wasp for 24h. Paralyzed larvae were removed and hemolymph samples were collected by cutting their abdominal prolegs with a pair of sterile scissors. Prior to hemolymph extraction, each larva was sterilized (70% ethanol) and rinsed with sterile water. 60µl of hemolymph collected per larva was mixed with 200µL saline (PBS). 20µl of this was placed on teflon coated diagnostic slide (8mm well) and fixed with 4% formaldehyde for 15 min to preserve the cell morphology, followed by two washes with PBT (PBS with 0.3% triton X). F-actin specific staining was done using alexafluor 488 (1:200, Thermo Fisher Scientific) for 2h. The sample was mounted with DAPI vectashield (Vector laboratories). Images were recorded on Olympus FV3000 confocal microscope using 20X objectives. Hemocytes with filapodia, pseudopodia or flaring (all indicators of spreading) were counted for comparative analysis (*n*=5). Hemocytes that retained intact shape and morphology were counted as viable while cells that showed blebbing or rupture were considered non-viable cells (Teng et al. 2016). Larvae not exposed to parasitoid served as control.

### Field Studies

To assess the potential of *B. brevicornis* in controlling *S. frugiperda*, a field experiment was conducted in FAW infested Maize field, Chikkaballapur District, Karnataka (13° 28′ N, 77° 72′ E, 918 m above sea level), India, during August–September 2019. The experiment was conducted in a Maize (cv. Rishi) field of 2000 m^2^, with a plant spacing of (75 × 20 cm) where plants were sown following ridges and furrow method in red sandy loam soil. Field inspection was done during early whorl stage to check for FAW infestation (20 days old) and the releases were made during true whorl stage of maize (25 days old) when FAW infestation reached its peak. The experimental layout was a randomized complete block design with two treatments (T1= control plot; T2= experimental plot where *B. brevicornis* was released). There were 20 replicates (block) per treatment (1000 m^2^). Each block had five plants. T1 was separated from T2 by a buffer plot (100 m) to reduce the probability of parasitoid moving to control plot. *B. brevicornis* was released on 25-days old crop infested with FAW. In total, three releases were made (at the rate of 4000 adults/ha) at weekly intervals. Parasitoids were released at ∼9.00 AM when temperature varied between (20-25 °C); humidity was 56-84% and wind speed was 14-24 km/h (Meteorological Centre, Bangalore, India) during all the three releases. 24 h old parasitoids were used for release in 1:1 ratio (male: female). They were transferred to plastic container covered with black muslin cloth. In the field, parasitoids were released at 5 spots by removing the muslin cloth and tapping the container while walking at a slow pace along the row. Pre-release larval population was also estimated in both the plots. Percent parasitism in larval population was estimated by counting the visible number of paralyzed/parasitized larvae after every release. To avoid destructive sampling in farmer’s field, only visible, live or paralyzed/parasitized larvae were counted. After three releases, to monitor the parasitoid activity in the field, five sentinel pouches with *Corcyra cephalonica* larvae (10 larvae/pouch) were introduced in T1 and T2 plots. These were collected post 24h to check parasitization.

### Statistical Analyses

To analyze the relationship between host ontogeny and parasitoid development, one-way ANOVA was performed considering instar stage (3^rd^, 4^th^ and 5^th^) as main effect and fecundity, larval hatching, percent pupae formation, adult emergence and percent female emergence as dependable variables respectively. To meet conditions of normality, data on fecundity and percent pupae formation were arcsine and square root transformed respectively. When ANOVA indicated a significant result (α < 0.05), all the relevant means were compared using Tukey’s post-hoc significance test at a level of 5%.

Data on total number of eggs laid per larva, percent emergence and fecundity were compared among the host instars (4^th^ and 5^th^) using un-paired *t*-test. Cell viability and cell spreading assay data were compared between control (un-paralyzed) and paralyzed larvae using un-paired *t* test. Field data on larval population were tested using independent *t*-test to compare the mean number of larvae/plant between released and control plot.

Percent parasitism on different host instars in cage study and percent parasitism by *B. brevicornis* in the field was analyzed using generalized linear model based on binomial distribution using logit link function. Data analysis was done using R statistical program (R core Team 2016) and SPSS Statistics version 25.0 (IBM). Assumptions of normality and homoscedasticity were tested before each test.

## RESULTS

### Performance and instar preference of *B. brevicornis* on two *Spodoptera* hosts

The lifetime fecundity of *B. brevicornis* was affected by the larval instar of *S. litura*. The highest fecundity was recorded on fifth instar larvae (253±17eggs, Suppl. video S1) while there was no significant difference between the third and fourth instar, which ranged between 138-173 (Table 1). There were no significant differences in percent larval hatching among the tested host instars (47-62%). However, highest percent pupa formation was observed on fifth instar (76%) compared to third and fourth instar. Percent adult emergence and the number of female emergence varied significantly; an average of 26 (±1) *B. brevicornis* females emerged from 5^th^ instar, which was highest among the instars tested. The female wasps survived for 19-26 days and host instar had no impact on their longevity (Table 1). The lifetime fecundity of *B. brevicornis* was highest on fifth instar *S. frugiperda* larvae as the female wasp deposited 293(±80) eggs (Table 2). Egg hatching percentage was lowest on third instar. Percent hatching was not significantly different between fourth and fifth instars (Table 2). 80% of the parasitoid larvae on the fifth instar larva formed pupae and adult emergence was also recorded highest on this host developmental stage (70%). The number of females emerged increased with host age, fifth instar recorded the highest number of females and the life span of these females was also high. Females emerged from fifth instar host larva had a life span of 24 days while females from fourth and third instar lived for 13 and 10 days respectively. Although percent hatching was not different between fourth and fifth instar, pupal formation, adult emergence, number of females and their life span was higher on fifth instar compared to fourth instar (Table 2). Third instar larvae recorded lowest on all these parameters among all the three developmental stages tested. Taken together, these results show that *B. brevicornis* development is dependent on host instar and 4-5^th^ host larval stage is most suitable in the case of both *Spodoptera* hosts.

**Table 1.** Biology of *Bracon brevicornis* on different instars of *Spodoptera litura*. Data represents mean ±SE, Different letters indicate statistical differences based on Tukey’s post hoc test after one-way ANOVA (*n* = 10).

| Larval instar | Fecundity<br>(mean ± SE) | % hatching<br>(mean ± SE) | % pupae<br>formation<br>(mean ± SE) | % adult<br>emergence<br>(mean ± SE) | No of<br>females<br>(mean ± SE) | Female life<br>span (days)<br>(mean ± SE) |
| --- | --- | --- | --- | --- | --- | --- |
| **3<sup>rd</sup>** | 138<sup>b</sup> ± 42 | 47 ± 1.5 | 50<sup>b</sup> ± 3 | 50<sup>b</sup> ± 1.3 | 13<sup>b</sup> ± 3 | 19 ± 4 |
| **4<sup>th</sup>** | 173<sup>b</sup> ± 32 | 53 ± 5 | 52<sup>b</sup> ± 5 | 52<sup>b</sup> ± 4 | 18<sup>b</sup> ± 3 | 24 ± 5 |
| **5<sup>th</sup>** | 253<sup>a</sup> ± 17 | 62 ± 8 | 76<sup>a</sup> ± 6 | 78<sup>a</sup> ± 9 | 26<sup>a</sup> ± 4 | 26 ± 1 |
| ***F*<sub>2, 27</sub>** | 8 | 3.2 | 24.7 | 25.5 | 9.4 | 3.5 |
| ***P*** | <0.001 | =0.06 | <0.0001 | <0.0001 | <0.0001 | =0.06 |

**Table 2.** Biology of *Bracon brevicornis* on different instars of *Spodoptera frugiperda*. Data represents mean ±SE, Different letters indicate statistical differences based on Tukey’s post hoc test after one-way ANOVA (*n* = 5).

| Larval instar | Fecundity<br>(mean ± SE) | % hatching<br>(mean ± SE) | % pupae<br>formation<br>(mean ± SE) | % adult<br>emergence<br>(mean ± SE) | No of<br>females<br>(mean ± SE) | Female life<br>span (days)<br>(mean ± SE) |
| --- | --- | --- | --- | --- | --- | --- |
| **3<sup>rd</sup>** | 108<sup>c</sup> ± 4.7 | 23<sup>b</sup> ± 1.5 | 25<sup>c</sup> ± 8 | 24<sup>c</sup> ± 4 | 11<sup>c</sup> ± 3 | 10<sup>b</sup> ± 0.8 |
| **4<sup>th</sup>** | 190<sup>b</sup> ± 43 | 47<sup>a</sup> ± 5.4 | 47<sup>b</sup> ± 5.3 | 45<sup>b</sup> ± 5.3 | 18<sup>b</sup> ± 5 | 13<sup>b</sup> ± 1.4 |
| **5<sup>th</sup>** | 293<sup>a</sup> ± 80 | 57<sup>a</sup> ± 8.5 | 80<sup>a</sup> ± 3.5 | 70<sup>a</sup> ± 6.2 | 33<sup>a</sup> ± 10 | 24<sup>a</sup> ± 5 |
| ***F*<sub>2, 12</sub>** | 5.48 | 9.8 | 41 | 28 | 13 | 8.1 |
| ***P*** | =0.002 | =0.003 | <0.0001 | <0.0001 | =0.002 | =0.005 |

In two-choice experiment in a petri dish, the female wasp always preferred to parasitize the fourth or fifth instar larvae against third instar of both the tested hosts. When the choice was between fourth and fifth instar *S. litura, B. brevicornis* chose fourth over fifth instar larvae (Figure S1a) (*P*<0.0001). In case of *S. frugiperda*, there was no preference for the same treatment (Figure S1b) (*P*=0.68).

### Impact of host plant, *Zea mays* on *B. brevicornis* interaction

When *S. litura* larvae on *Zea mays* were presented to *B. brevicornis* in a cage, mimicking natural conditions, their parasitism rate was 8 (±5), 72 (±5.2) and 76 (±7.4) % against 3^rd^, 4^th^ and 5^th^ instar respectively (Figure 1a). *B. brevicornis* showed higher parasitism rate on 4^th^ and 5^th^ instar *S. litura* larvae similar to results obtained in earlier choice assay in the absence of the plant. The performance on *S. frugiperda* showed similar results, where 16 (±4), 76 (±4) and 84 (±5.6) % parasitism was recorded on 3^rd^, 4^th^ and 5^th^ instar, respectively (Figure 1b) (*P*<0.0001). The average number of parasitoid eggs laid on fourth and fifth instar of *S. litura* larvae was 5.9 (±1.3) and 5.5 (±1.2), respectively. These parasitized larvae with eggs were incubated at laboratory conditions. From these eggs, 70% of adults could emerge from fourth instar while fifth instar supported 66% of adult emergence from *S. litura* (Figure 2a, b). Although statistically there was no significant difference between fourth and fifth instar larvae of *S. litura* in terms of *B. brevicornis* oviposition and adult emergence, the adult longevity was higher from fifth instar host (Figure 2c, *P*<0.0009).

**Figure 1:**
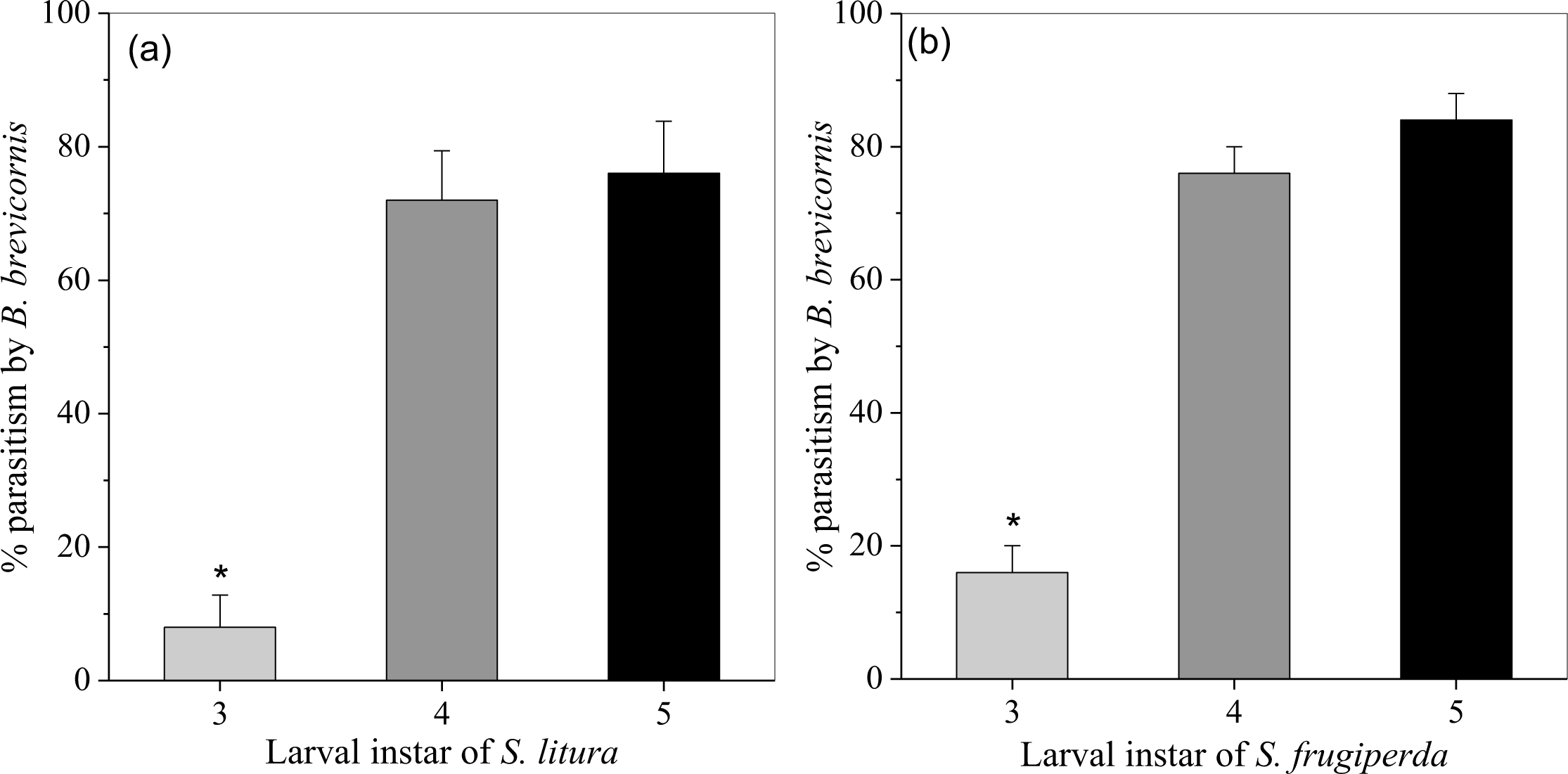
Percent parasitism by *B. brevicornis* in the presence of host plant against (a) *S. litura* and (b) *S. frugiperda* in cage study. Data represents mean ±SE, statistical difference is based on generalized linear model (*n* = 5). 3: third, 4: fourth and 5: fifth instar larvae.

**Figure 2:**
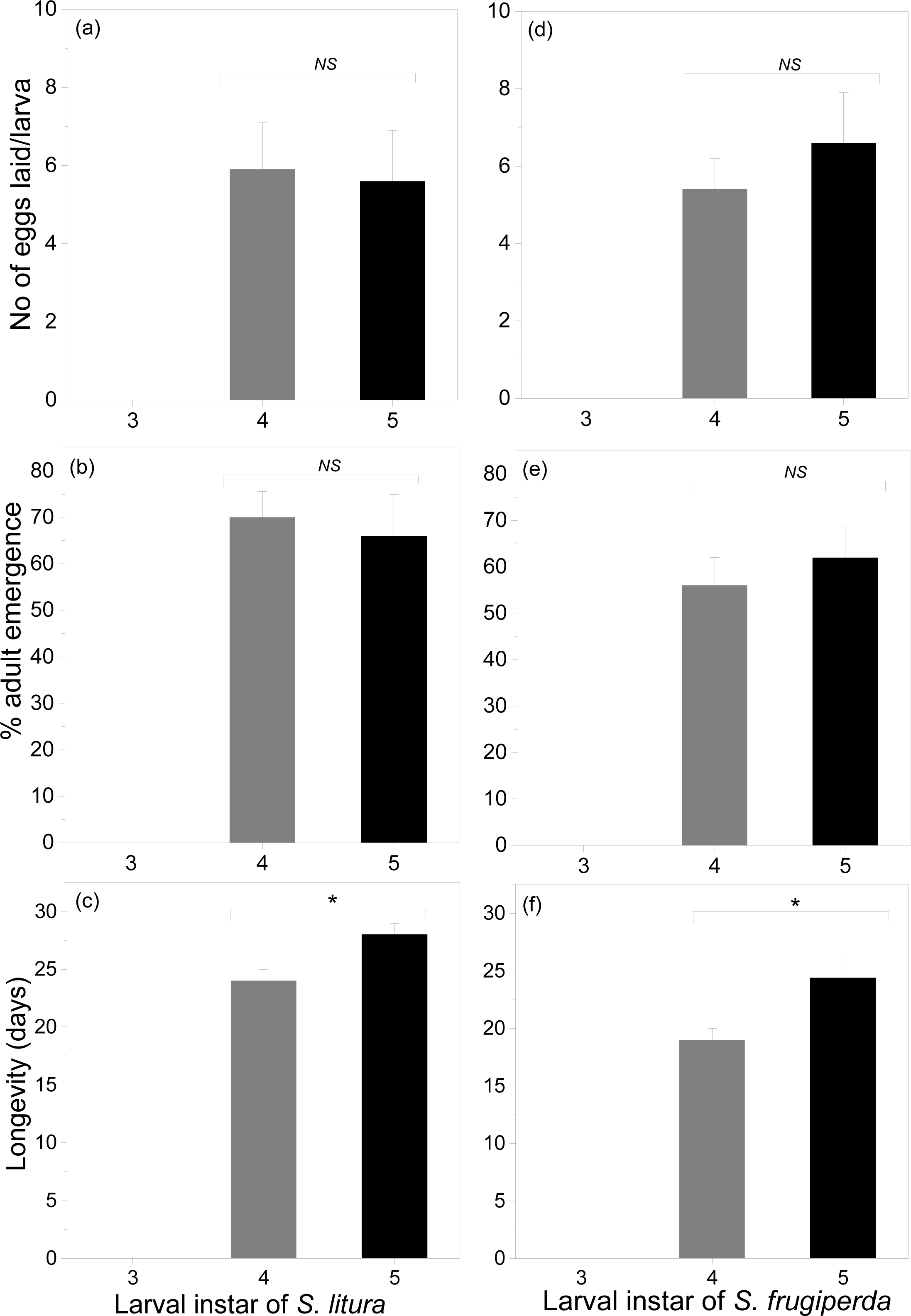
Biology of *B. brevicornis* in the presence of host plant in cage study. No of eggs laid per larva, percent adult emergence and longevity of the adult emerged from parasitized (a, b, c) *S. litura* and (d, e, f) *S. frugiperda* larval instars. (Data represents mean ±SE, statistical differences are based on *t*-test (*n* = 10). 3: third, 4: fourth and 5: fifth instar larvae.

In case of *S. frugiperda*, the female parasitoid deposited 5.4 (±0.8) on fourth and 6.6 (±1.3) number of eggs on fifth instar larvae (Figure 2d). In laboratory conditions, the adult emergence rate from fourth and fifth instar ranged from 57-62%. (Figure 2e). Our results show that *B. brevicornis* can perform equally well on fourth and fifth instar of larvae of the hosts. However, the longevity of the emerged adults from fifth instar was significantly higher (Figure 2f, *P*<0.0001).

### Alteration in host immune-competence

Total hemocyte numbers were less in paralyzed larvae compared to control. In paralyzed larvae, the viability and spreading capacity of hemocytes was significantly reduced (Figure 3a, c) in case of *S. litura* (*P*<0.0001). Staining with phalloidin-DAPI showed these hemocytes had membrane blebbing and ruptured structure due to which they lost their spreading behavior (Figure 3e, f). Parasitoid-induced immune suppression did not differ significantly from *S. litura* in case of *S. frugiperda* larvae. The viable hemocytes number was reduced after paralysis by *B. brevicornis* and these hemocytes rapidly lost their shape and spreading ability (Figure 3b, g). Staining with phalloidin-DAPI showed similar morphology as in the case of *S. litura* (Figure 3d, h). These results indicate that host regulation by *B. brevicornis* affects cellular immunity of host larvae and does not differ significantly between the two *Spodoptera* sp. tested.

**Figure 3:**
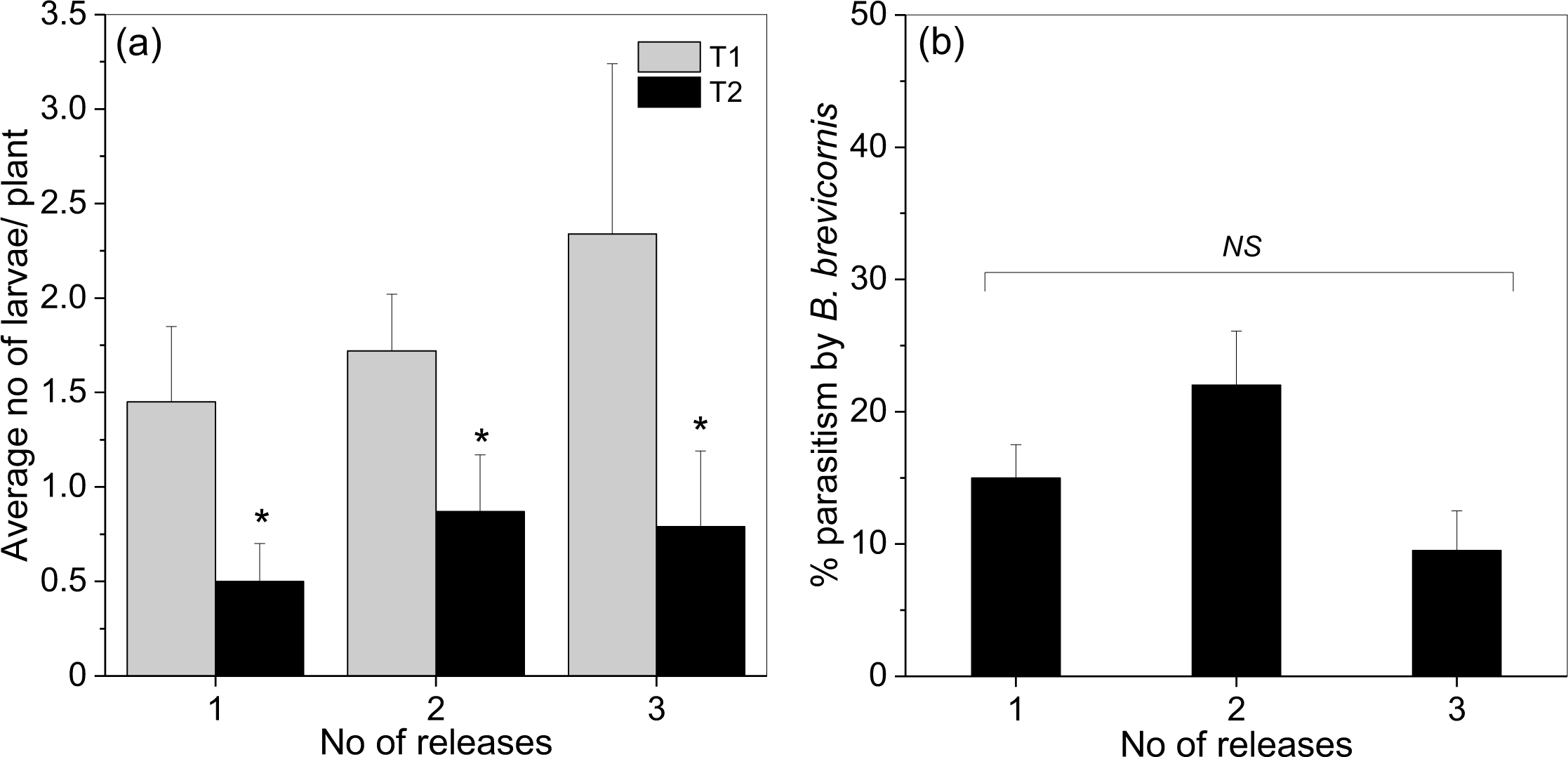
Regulation of host immune-competence by *B. brevicornis*. Changes in hemocyte viability and spreading post paralysis of (a, b) *S. litura and* (c, d) *S. frugiperda*. Phalloidin-DAPI stained hemocytes in control and paralyzed larvae of (e) *S. litura* and (g) *S. frugiperda*. Alteration in hemocyte spreading activity in control and paralyzed larvae of (f) *S. litura* and (h) *S. frugiperda*. Data represents mean ±SE, statistical differences are based on *t* test (*n* = 5), bars: 10µm.

### From lab to field: *B. brevicornis* performance on FAW under field conditions

To test the potential of *B. brevicornis* in pest management, the wasps were released in a FAW infested maize field (Fig. S2, suppl. video S3). *Bracon brevicornis* were released three times at weekly intervals and the average number of larvae recorded per plant before and after each release was significantly reduced compared to control plot (T1) (*P*<0.0001) (Figure 4a). This led to a total larval reduction of 65, 47 and 50% post first, second and third release, respectively. Percent parasitism by *B. brevicornis* in treatment plot ranged between 10-22% after releases. There was no significant difference between different releases (Figure 4b). After the releases, the remaining live larvae were noted to be largely in early developmental stages (1^st^, 2^nd^ and 3^rd^ instar), which is possibly be due to fresh egg laying by moths coming from the neighboring fields or larvae which escaped from parasitization and could complete their life cycle (Figure S2). To check if the parasitoids released reached control plots, sentinel pouches containing *C. cephalonica* larvae were kept in both plots after our experiment. No parasitization was recorded in control plot (T1) while in the experimental plot (T2), 40% of *C. cephalonica* larvae were parasitized by *B. brevicornis*.

**Figure 4:**
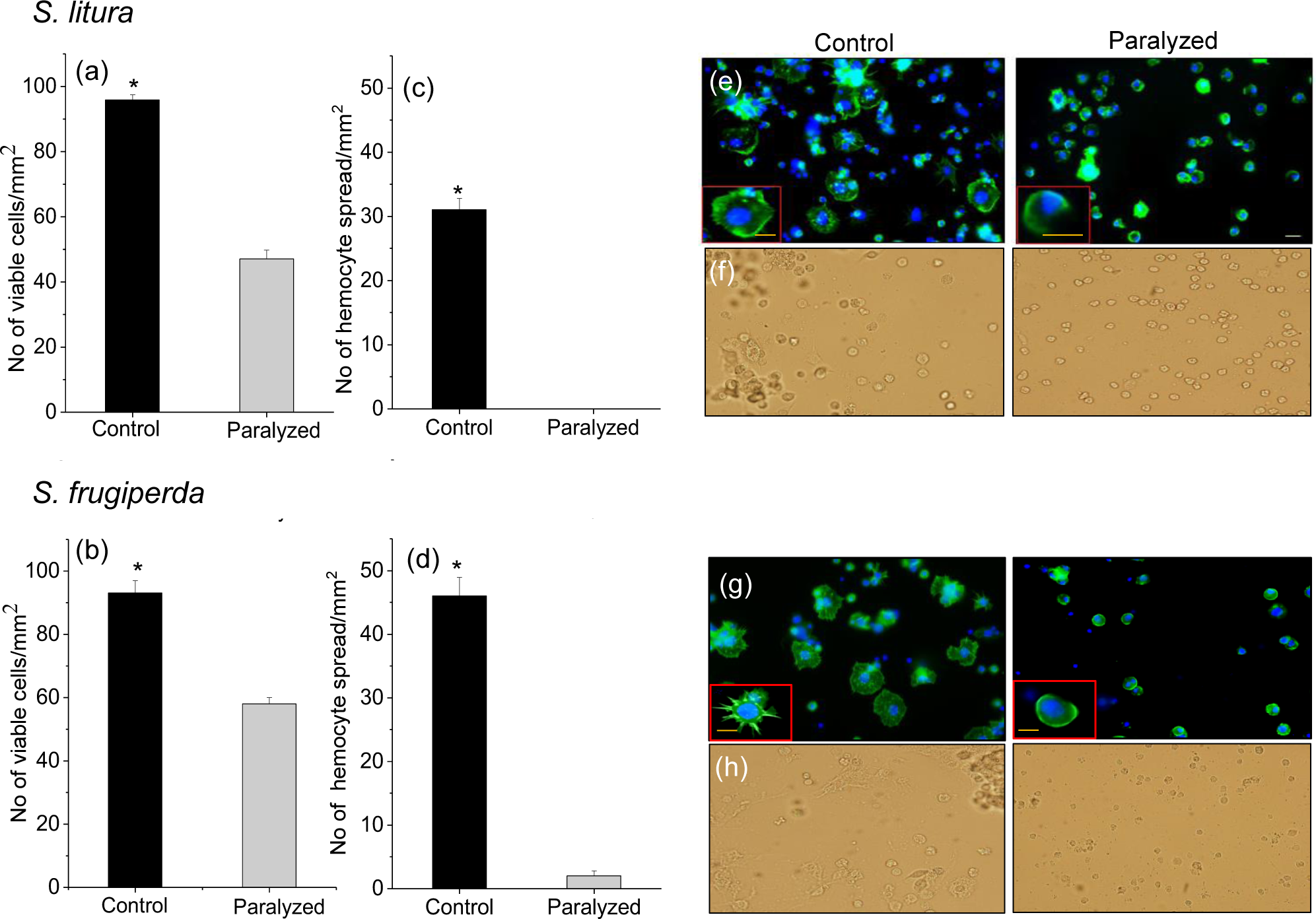
The effectiveness of *B. brevicornis* in FAW infested maize field. (a) average number of larvae/plant after each release in T1 (Control plot) and T2 (experimental plot) (b) Percent parasitism by *B. brevicornis* in experimental plot. Data represents mean ±SE, statistical significance is based on t test and generalised linear model analysis (*n* = 20).

## Discussion

Appropriate host selection is crucial for parasitoid progeny development (Mattiacci and Dicke 1995) and host developmental stage influences parasitoid attack rate, survival and sex ratio (Jervis et al. 2007; Kant et al. 2012; Li et al. 2006). Here, using *S. litura* and *S. frugiperda*, we examined the performance of *B. brevicornis* on different host developmental stages. It is interesting to note that three larval instars (3-5^th^) of both hosts were susceptible to *B. brevicornis* attack and the parasitoid could complete its lifecycle on 3-5^th^ instar of these two hosts in the laboratory. However, the performance of the parasitoid in terms of average number of eggs laid, adult emergence and lifespan was best on 5^th^ followed by 4^th^ instar, which indicates that late larval stages of these *Spodoptera* hosts are most suitable for *B. brevicornis* development. Our results are in accordance with Malesios and Prophetou-Athanasiadou, (2014), where biology of *B. brevicornis* on *Plodia interpunctalla* (Hübner) was shown to be strongly affected by host larval instar and preference of late instars was reported. Host suitability depends on factors like oviposition attempts, nutrient content and successful progeny development from them (Wiedemann et al. 1992). *B. brevicornis* is a gregarious parasitoid and their developing larvae acquire nutrients by constant feeding of host hemolymph (Suppl. video S2). Hence, the host must serve as an adequate nutrient pool to support more than one developing parasitoid larvae until they reach pupal stage. As 4-5^th^ instar larvae contain higher amount of hemolymph (approx. 60ul), they probably serve as better hosts. Although *B. brevicornis* could attack and parasitize all instars tested, the performance was best on late instar and choice test done clearly indicated that *B. brevicornis* prefers older instar larvae of both the hosts. Whether *B. brevicornis* uses odors or other cues to distinguish larval instar remains to be studied.

Parasitoids like *Cotesia flavipes* (Cameron) also show similar preference towards late larval instars of the host *Diatraea saccharalis* (F.) (Wiedenmann et al. 1992), while *Cotesia rubecula* Marshall prefers to oviposit in early first larval instars of *P. brassicae* (L.) and *P. napi* (L.) (Brodeur and Geervliet 1995). Analysis by Brodeur and Vet (1995) shows that often endo-parasitoid oviposit in the early instars to avoid the well-developed immune system of the host. In the case of *S. litura*, previous study shows that there is an increase in immune response with ontogeny (Ghosh and Venkatesan 2019). However, the immune status did not influence the behavior of *B. brevicornis* as they could down-regulate cellular immunity of both hosts at 4^th^ and 5^th^ instar stages. Since late larval stages would serve as better nutrient resources and parasitoid-derived factors could successfully suppress host cellular immunity, *B. brevicornis* performance was higher on 4^th^ and 5^th^ instar and the female wasp showed a preference for late larval hosts in two-choice assay. Differences in the biochemical status, immune-competence, humoral immune responses and/or odor of host larvae could contribute towards parasitoid preference.

Further, we tested the impact of host instar on parasitoid progeny. Since female offspring are more important for reproductive output (Godfray 1994) and for further biological applications, we assessed the sex ratio on different host instars. Similar to earlier results, 4^th^ or 5^th^ instar host larvae supported more female parasitoids. Taken together, our findings show that selection of late larval instar hosts increases the overall fitness of *B. brevicornis*. Next, we evaluated if the parasitoid interaction is altered in the presence of host plant, *Zea mays*. Previous studies showed that host plant on which the herbivorous larvae develop can significantly alter parasitoid behavior (Li et al. 2014; Gols et al. 2008; Sarfraz et al. 2019; Stoepler et al. 2011). Plants are known to release specific bouquet of volatiles upon herbivory that function as attractive cues for parasitoids. The role of herbivore induced plant volatiles (HIPV) in mediating parasitoid interactions is well documented. For example, females of *Cotesia vestalis* (Haliday) are attracted only by HIPV from *Brassica rapa* plants infested with *Plutella xylostella* (Linnaeus) (Kugimiya et al. 2010). Such HIPV also help parasitoids to distinguish between larval stages, plant species and different levels of damage (Girling et al. 2011). These studies show that host plant chemistry can impact parasitoid host choice. In our study, *B. brevicornis* showed similar preference for older instars in the presence and absence of host plant. Interestingly, in test tube assay, 3^rd^ instar could support parasitoid egg hatching and subsequent development, however, on host plants, *B. brevicornis* showed very low parasitism rate and did not oviposit or develop on 3^rd^ instar larvae of both the hosts. This could be due to pre-oviposition interactions between host and parasitoid. We observed that, early instar larvae are comparatively more agile and can also hide well inside the maize whorl, which could make the oviposition experience difficult for *B. brevicornis*. This is in line with previous study where parasitoid was reported to use larval cues to discriminate hosts and avoid hosts that counterattack by thrashing, biting, and regurgitating (Feltwell 1982). The role of plant volatiles induced by different larval instars on parasitoid choice warrants further investigation.

Encouraged by lab-based study, we evaluated the performance of *B. brevicornis* in a FAW infested maize field. Our results show that *B. brevicornis* can efficiently reduce the infestation and parasitize FAW larvae even under field conditions. Our first release showed a higher reduction in average larval population per plant than control compared to second and third release due to the presence of more late-instar larval population (4^th^ and 5^th^ instars). Since no other pest control was implemented in the experimental plot, factors such as fresh egg laying by moths from existing or neighboring field, presence of younger instar host larvae (during second and third release) and/or un-parasitized larvae could be possible reasons for less percent reduction in larval population during other releases compared to the first. However, the significant difference between control and experimental plot after release is interesting and indicates that *B. brevicornis* could be used in augmentative control of FAW. Also, the average larvae per plant reduced significantly after three releases. The visible parasitism rate of in the field ranged from 10-22% in line with previous studies on *Bracon hebetor* and black headed caterpillar, where 8-27% parasitism rate was reported in coconut palm field (Rao et al. 2018). Since we counted only visible larvae, it is possible that our sampling method could be an underestimation of parasitism rate. Additionally, overlapping generations of FAW were found in the field that could lead to longer search time by the parasitoids.

Taken together, *B. brevicornis* can be employed in combination with egg parasitoids like *Telenomus remus* (Kenis et al. 2019) and *Trichogramma* spp. to obtain a greater success in pest management. Interestingly, *B. brevicornis* feeds on host hemolymph prior to egg laying. They also fed on 3^rd^ instar larvae post paralysis, thus killing it. Hence, we anticipate that they might be able to control 3^rd^ instar larvae also. These results make *B. brevicornis* a potential candidate for biological control program of these hosts in real field conditions. Furthermore, *B. brevicornis* is a generalist parasitoid and can produce a very high number of progeny on various lepidopteran pests like *Corcyra cephalonica* (Dabhi et al. 2011), therefore, mass production can easily be done by insectaries. In summary, the current study shows that *B. brevicornis* can be an effective biological control agent against *S. frugiperda* and *S. litura* larvae. Our findings provide valuable information for the development of augmentative biological control program and IPM strategies against invasive pest *S. frugiperda*.

## Author Contributions

EG and RV designed and carried out experiments, performed field work, analyzed data and drafted the manuscript. RV conceptualized, supervised the project and experimental design, performed field work, acquired resources and drafted the manuscript. All authors read and approved the final manuscript.

## Acknowledgments

Funding from NCBS, Max Planck Gesellschaft (Partner Group Program), DBT-NER grant and Ramanujan Fellowship is acknowledged. The authors thank Indian Council of Agricultural Research, New Delhi for research support and Dr. Sane lab, NCBS for parasitoid behavior recording. The authors thank TNAU for initial culture of *B. brevicornis*.

## Legends

Figure S1: Effect of host larval instar on *B. brevicornis* oviposition choice. Two-choice test between third vs fourth, third vs fifth and fourth vs fifth instar larvae of (a) *S. litura*, (b) *S. frugiperda* presented to one female parasitoid. Statistical significance is based on un-paired *t* test (*n* = 5).

Figure S2: Stages of *S. frugiperda* found in maize field (a) female moth, (b) freshly laid eggs, (c) newly hatched larvae, (d) early instar of larva, (e, f) late instar of larvae, (g, h) paralyzed larvae post *B. brevicornis* release, (i) *B. brevicornis* larvae on *S. frugiperda*.

Video S1: Oviposition behaviour of *B. brevicornis* on paralyzed *S. litura* larvae.

Video S2: *B. brevicornis* larvae feeding on paralyzed *S. litura* larvae.

Video S3: Interaction between *B. brevicornis* female and *S. frugiperda* larvae in maize field.

