## Supplementary figures and images for "Performance of *Bracon brevicornis* (Wesmael) on two *Spodoptera* species and application as potential biocontrol agent against fall armyworm"

### Supplemental Figures

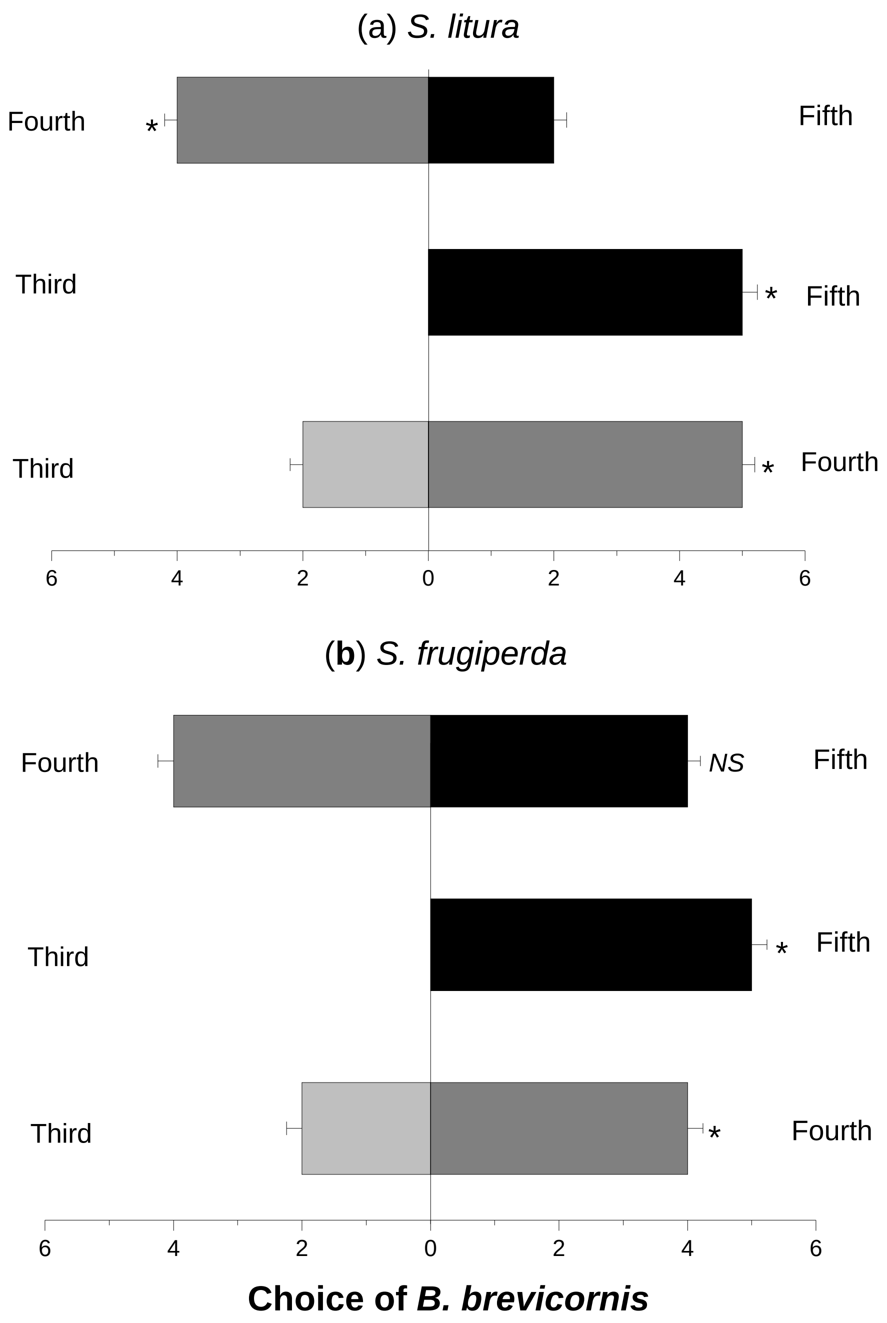

Figure S2

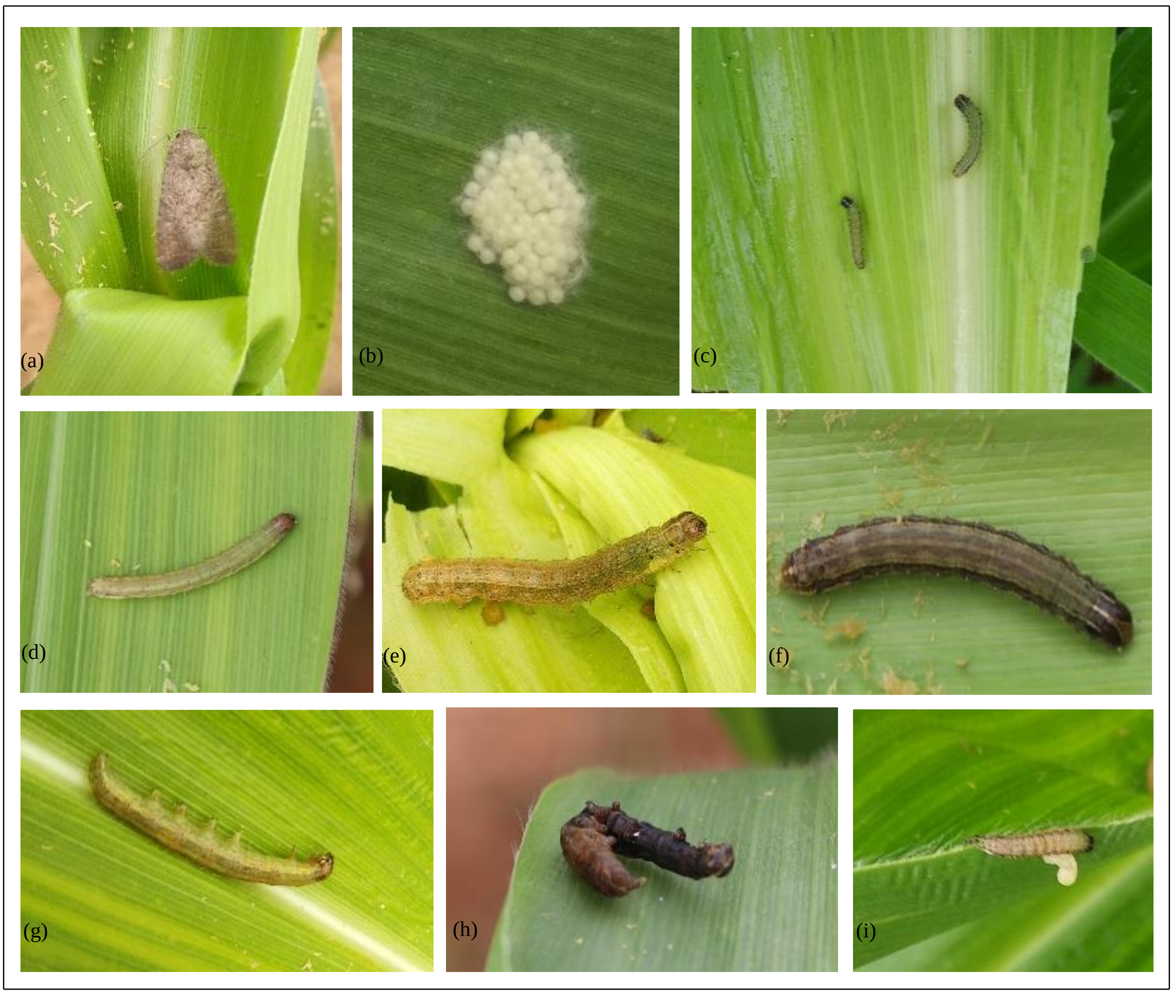
